# Structure of human aldehyde oxidase under tris(2-carboxyethyl)phosphine-reducing conditions

**DOI:** 10.64898/2026.03.25.713928

**Authors:** Carolina Videira, Mariam Esmaeeli, Silke Leimkühler, Maria João Romão, Cristiano Mota

## Abstract

The importance of human aldehyde oxidase (hAOX1) has increased over the last decades due to its involvement in drug metabolism. Inhibition studies concerning hAOX1 are extensive and a common reducing agent, dithiothreitol (DTT), was recently found to inactivate the enzyme. However, in previous crystallographic studies of hAOX1, DTT was found to be essential for crystallization. To surpass this concern another reducing agent used in crystallization trials. Using tris(2-carboxyethyl)phosphine (TCEP), a sulphur-free reducing agent, it was possible to obtain well-ordered crystals from hAOX1 wild type and variant, hAOX1_6A, which diffracted beyond 2.3 Å. Instead of the typical star-shaped crystals of hAOX1, at pH 4.7, plates are obtained in the orthorhombic space group (P22_1_2_1_) with two molecules in the asymmetric unit. Activity assays with the enzyme incubated with both reducing agents show that contrary to DTT, TCEP does not lead to irreversible inactivation of the enzyme. The replacement of DTT with TCEP in crystallization of hAOX1 provides a strategy to circumvent enzyme inactivation during crystallographic studies, allowing future applications of new assays, such as time-resolved crystallography.

## 1. Introduction

Aldehyde oxidases (AOXs; EC 1.2.3.1) are cytosolic molybdoflavoproteins that belong to the Xanthine Oxidoreductase (XOR) family (Garattini *et al*., 2008) and require a molybdenum cofactor (Moco), two [2Fe-2S] clusters and a flavin adenine dinucleotide (FAD) to be catalytically active (Garattini *et al*., 2003). AOXs are active as homodimers with each 150 kDa monomer comprising 3 functional domains: the N-terminal domain harbouring two [2Fe-2S] centers (20 kDa), followed by the central FAD domain (40 kDa) and, finally, the catalytic Moco domain (90 kDa) where the substrate pocket is located (Coelho *et al*., 2015).

Humans only express one active form of AOX, assigned hAOX1, while mice have four isoforms (Garattini *et al*., 2008). Although widely expressed throughout the organism, the higher expression levels are found on adipose tissue, adrenal glands and liver (Terao *et al*., 2016). hAOX1 is relevant in drug metabolism, specifically during phase I and activation of prodrugs (Pryde *et al*., 2010; Mota *et al*., 2018; Guo & Lerner-Tung; Clarke *et al*., 1995), and some of its substrates are intermediates resulting from cytochrome P450 metabolism (Garattini & Terao, 2012; Romão *et al*., 2017). The hAOX1 involvement in this pathway has been investigated in the past years, with an increasing interest, especially for pharmaceutical companies in drug design campaigns (Mota *et al*., 2018). The physiological role and potential implications of hAOX1 in disease are however still poorly understood (Mota *et al*., 2018), but some studies have unveiled its possible involvement in fat-related metabolism (Weigert *et al*., 2008; Heid *et al*., 2020; Buechler *et al*., 2002; Sigruener *et al*., 2007).

hAOX1 catalyzes oxidation of N-heterocycles and aldehydes, reduction of N/S-oxides and nitro derivatives and hydrolysis of amide bonds (Terao *et al*., 2016; Hartmann *et al*., 2012). While the related XOR has a well-established role in the final steps of purine catabolism (Moriwaki *et al*., 1997), hAOX1 is a promiscuous enzyme, a feature associated with a wider substrate pocket that allows the accommodation of diverse and bulky molecules (Mota *et al*., 2018). Accurate prediction of novel substrates and the pharmacokinetic behaviour of hAOX1 remains challenging, and additional structural and functional data are required to elucidate the broad substrate specify of the enzyme.

The inhibition of hAOX1 has been extensively investigated and is well-characterised *in vitro* (Obach *et al*., 2004; Esmaeeli *et al*., 2023, Mota *et al*., 2021; Coelho *et al*., 2015). In particular, the capacity of dithiothreitol (DTT) to inactivate hAOX1 has been demonstrated (Esmaeeli *et al*., 2023). Cyanide and DTT are proposed to inhibit the enzyme through a similar mechanism, by removing the sulfido ligand of the Mo atom, which is crucial for the enzyme’s activity (Esmaeeli *et al*., 2023; Massey & Edmondson, 1970).

The previously reported crystallization conditions for hAOX1 required a pre-incubation step with DTT (Coelho *et al*., 2015; Mota *et al*., 2021), which should impairs unspecific intermolecular interactions particularly between cysteines on the surface of the enzyme, and allow crystal formation. Nevertheless, since DTT inactivates the enzyme, it is expected that it may compromise structural studies related to the enzyme’s activity.

Here, we report a new crystallization condition using an alternative reducing agent, tris(2-carboxyethyl)phosphine (TCEP), to obtain hAOX1 crystals. The activity assays indicated that contrary to treatment with DTT, the enzyme is not inactivated in the presence of TCEP.

## 2. Material and Methods

### 2.1. Mutagenesis, expression and purification of hAOX1

Plasmid pTrc_His_AOX1_6A encoding the hAOX1_6A variant (C161A/C165A/C170A/C171A/C179A/C180A) was generated by site-directed mutagenesis using the primers listed in Table 1 (Esmaeeli, PhD thesis, 2022; Zheng et al., 2004), with vector pTHcohAOX1as the template (Foti et al., 2016). *E. coli* TP1000 cells (Palmer *et al*., 1996) were transformed by Heat Shock with the vectors (Hanahan, 1983) and protein production was performed as previously described (Foti *et al*., 2016; Esmaeeli *et al*., 2023). After purification, the protein was concentrated to 10 mg/mL in 50 mM Tris-HCl pH 8, 200 mM NaCl, 1 mM EDTA and stored at -80 ºC.

**Table 1.**
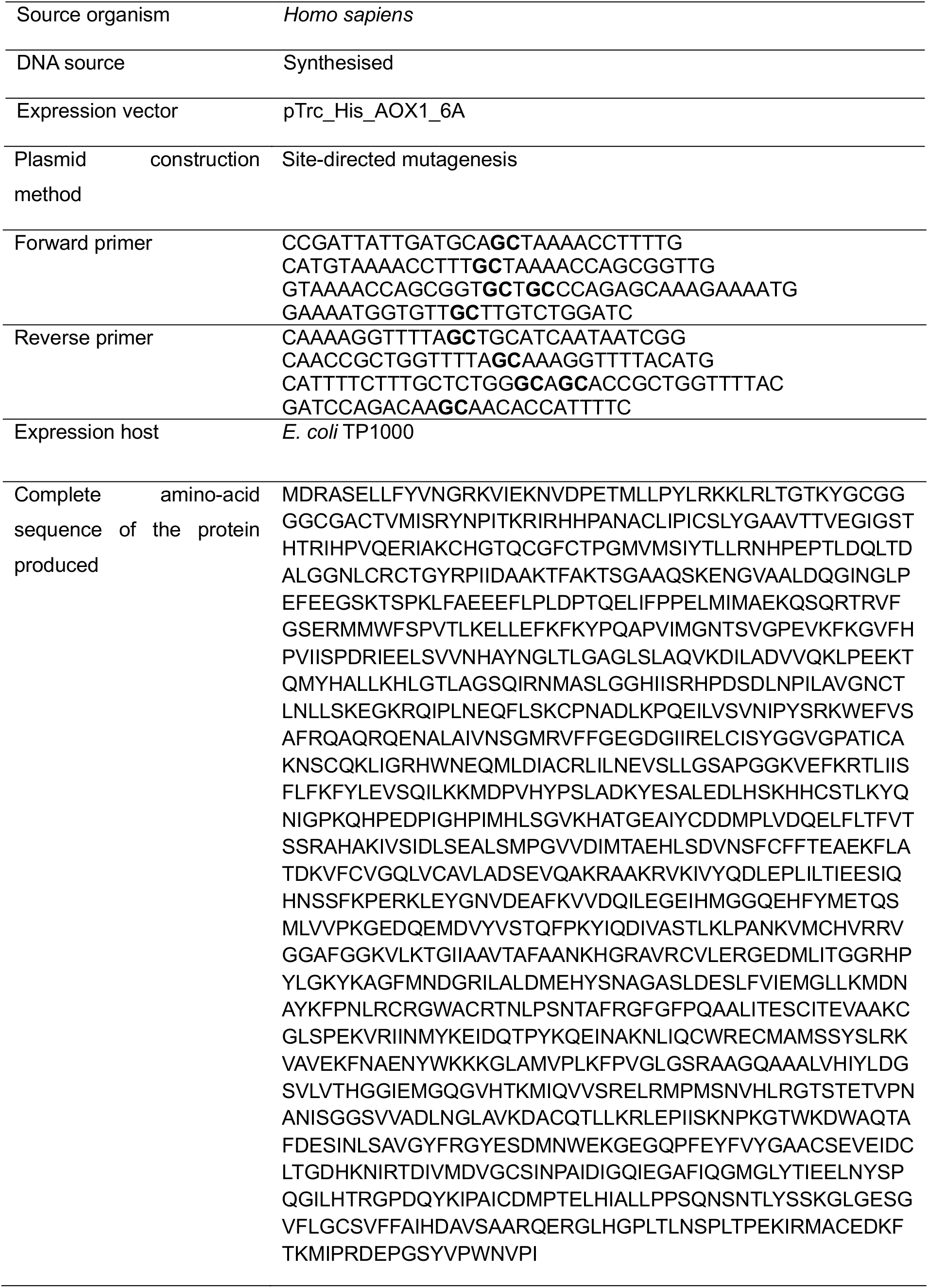
Macromolecule production information.

### 2.2. Crystallization, data collection, structure solution and refinement

Prior to crystallization, protein samples were incubated for at least 10 minutes with 1 mM tris(2-carboxyethyl)phosphine (TCEP; Sigma-Aldrich). Crystallization trials were performed at 4 °C and 20 °C using the hanging-drop vapor-diffusion method in 24-well plates XRL plate (Molecular Dimensions). Initial screening matrices were prepared by varying poly(ethylene glycol) 3350 (PEG3350; Sigma-Aldrich) concentration between 8% and 22% (w/v) in 2% increments in the presence of 0.1 M sodium malonate (Sigma-Aldrich) adjusted to different pH values (pH 4.5, 4.7, 5.0, 5.1, 5.3, 5.5, or 6). Two different protein concentrations of 5 and 10 mg/mL were tested.

Hanging drops consisted of 4 μL of protein solution (5-10 mg/mL in 50 mM Tris-HCl pH 8, 200 mM NaCl, 1 mM EDTA supplemented with 1mM TCEP) mixed with 2 μL of precipitant solution, and were equilibrated against 500 μL of reservoir solution containing 0.1 M sodium malonate and 10% PEG3350, at 4 ºC, as previously described (Mota *et al*., 2021). The best crystals of hAOX_6A appeared within 48h and grew for one week to a max size of 0.1 mm (Fig. 2) with the condition consisting of 12% PEG 3350, 100 mM sodium malonate pH 4.7. To stabilize the crystals, 4 μL of harvesting solution (100 mM sodium malonate pH 4.7 and 12% PEG3350) was added to the crystallization drop, and one crystal was transferred to a cryoprotectant solution (harvesting solution supplemented with 20% (v/v) glycerol) and flash-cooled in liquid nitrogen. X-ray diffraction data were collected at 100 K at beamline ID30A-3 (von Stetten *et al*., 2020), at the European Synchrotron Radiation Facility (ESRF; Grenoble, France). Data processing was done with STARANISO (Tickle *et al*., 2016). The structure was solved by MR with Phaser ((IUCr) Phaser crystallographic software), using the structure of the wild-type (wt) hAOX1-free form (PDB: 4UHW) as the search model. Manual model building, as well as iterative refinement, were carried out using Coot (Emsley *et al*., 2010, version 0.9.8.93), REFMAC (Murshudov *et al*., 2011, version 5.8.0425), and PDB REDO (Joosten *et al*., 2009). Stereochemical restraints were generated using phenix.elbow (Moriarty *et al*., 2009, version 1.20.1-4487) The quality of the model was assessed using Iris and MolProbity (Davis *et al*., 2007, Rochira *et al*., 2021). All images were created and obtained with PyMOL (Schrödinger, Versions 2.5.3 and 3.1.6.1). Information regarding crystallization conditions, data collection, processing and refinement statistics are available in Tables 2, 3 and 4 respectively.

**Table 2.**
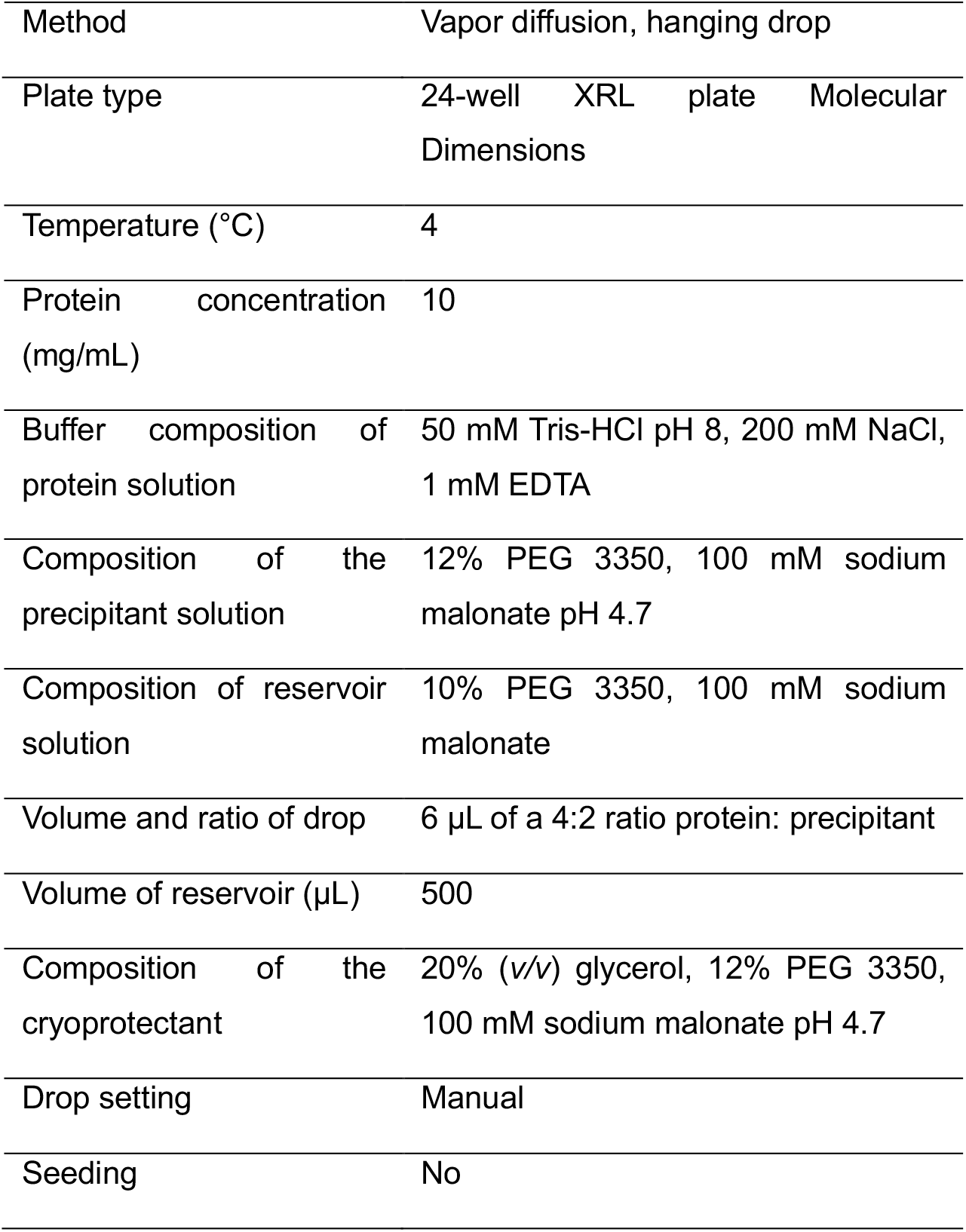
Summary of Crystallization conditions.

**Table 3.** Data collection and processing statistics. Values given in parentheses are for the highest resolution shell.

|  |  |
| --- | --- |
|  | hAOX1_6A + TCEP STARANISO |
| PDB code | 28MB |
| Diffraction source | ID30A-3, ESRF |
| Wavelength (Å) | 0.96770 |
| Temperature (K) | 100 |
| Detector | Eiger1 X 4M |
| Crystal to detector distance (mm) | 197.34 |
| Total rotation range (°) | 180 |
| Exposure time per degree (s) | 0.032 |
| Exposure time per image (s) | 0.0032 |
| Space group | P2<sub>1</sub>2<sub>1</sub>2 |
| a, b, c (Å) | 191.1, 273.0, 77.8 |
| Resolution range (Å) | 95.6-2.3 (2.6-2.3) |
| Anisotropic high-resolution limits (Å) | 3.0, 3.0, 2.3 |
| Total no. of reflections | 708883 (36291) |
| No. of unique reflections | 100292 (5015) |
| Completeness (overall/inner shell/outer shell) (%) |  |
| (ellipsoidal) | 94.5/ 99.9/81.4 |
| (spherical) | 55.7/ 99.9/8.8 |
| Redundancy | 7.1 (7.2) |
| (I/σ (I)) | 6.6 (1.6) |
| CC<sub>1/2</sub> | 100 (50) |
| R<sub>meas</sub> | 0.350 (1.554) |

**Table 4.** Refinement statistics. Values given in parentheses are for the highest resolution shell.

|  |  |
| --- | --- |
|  | hAOX1_6A + TCEP |
| PDB code | 28MB |
| Resolution range (Å) | 95.6-2.3 (2.6 - 2.3) |
| Final *R*<sub>cryst</sub> | 0.210 (0.347) |
| Final *R*<sub>free</sub> | 0.241 (0.404) |
| No. of non-H atoms | 20293 |
| Protein | 20164 |
| Ligand | 87 |
| Ion | 13 |
| Water | 29 |
| R.m.s deviations |  |
| Bonds (Å) | 0.007 |
| Angles (°) | 1.27 |
| Average *B*-factors<br>(Å<sup>2</sup>) |  |
| Protein | 31.3 |
| Ramachandran Plot |  |
| Most favoured (%) | 96.7 |
| Outliers (%) | 0.27 |
| Molprobity score | 1.48 |
| Clash Score | 2.35 |

### 2.3. hAOX1 activity assays

Kinetic assays to evaluate the effect of reducing agents on hAOX1 activity were conducted based on previous studies from (Esmaeeli *et al*., 2023). The wt hAOX1 (10 μM) was incubated with either DTT or TCEP at a 100-fold molar excess (1 mM final concentration) in 50 mM Tris-HCl, 200 mM NaCl, 1mM EDTA, pH 8.0, in a total reaction volume of 20 μL. A control sample containing only the enzyme in buffer was included. Incubations were carried out for 30 min, 4 hours and overnight (O/N). After the O/N incubation, the samples were subjected to buffer exchange using a 10 K centrifugal filter unit (Milipore®) to remove the reducing agents. Activity assays were performed in triplicate, using molecular oxygen as the electron acceptor in a final reaction volume of 500 μL. Each reaction contained 40 μM of the substrate phenanthridine (Sigma-Aldrich), in assay buffer (50 mM Tris-HCl, 200 mM NaCl, 1mM EDTA, pH 8.0). Reactions were initiated by the addition of enzyme (100 nM final concentration). Enzymatic activity was monitored by measuring the formation of phenanthridinone at 321 nm (ε321=6400 M^-1^ cm^-1^) for 300 s using an Evolution 201 UV-Visible Spectrophotometer (ThermoFisher Scientific).

## 3. Results and discussion

### 3.1. Crystal structure of hAOX1_6A in the presence of TCEP

To circumvent the use of DTT in the crystallization conditions, a variant of hAOX1 was designed by site-directed mutagenesis by mutating to alanines 6 cysteines (C161A/C165A/C170A/C171A/C179A/C180A) present at the protein surface in a 71-residue solvent-exposed and flexible loop (A160**C**KTF**C**KTSG**CC**QSKENGV**CC**…SQ231) (Esmaeeli, PhD thesis, 2022). Despite multiple efforts and extensive screenings, the variant hAOX1_6A could never be crystallized in the absence of a reducing agent.

A common substitute of DTT is Tris(2-carboxyethyl)phosphine (TCEP), with advantages over DTT, namely its stability to oxidation and the absence of thiol groups in its structure. A new crystallization setup was performed to optimize conditions and crystallize hAOX1 (both wt and variant 6A) in the presence of TCEP. To reduce the substantial protein precipitation observed in drops, different protein concentrations (either 5 mg/mL or 10 mg/mL) were used. Nevertheless, drops with lower protein concentration either led to the formation of smaller crystals or crystals with a non-regular morphology. Therefore, a protein stock solution at 10 mg/mL was used for the preparation of crystallization drops.

Typical DTT-grown hAOX1 crystals have a star-shaped morphology (Fig. 1, panel A) but are often multiple and diffract up to 2.8 Å (Coelho *et al*., 2015).When TCEP is used to crystalize hAOX1, both wt and 6A variant grow in the form of plate crystals under different PEG concentrations at pH 4.7, in both 4ºC and 20 °C (Fig. 1, panel B). Nevertheless, at higher pH, star-shaped crystals are obtained for both wt and variant. The best crystals were obtained for hAOX1_6A, and a single crystal, obtained at 12% PEG 3350, 0.1 M sodium malonate pH 4.7 in the presence of 1 mM of TCEP at 4 °C diffracted up to 2.3 Å in ID30A-3 beamline (von Stetten *et al*., 2020). This was an improvement since crystals from the wt protein always diffracted poorly (up to 2.8 Å) (Coelho *et al*., 2015; Mota *et al*., 2019).The diffraction data were processed in the orthorhombic space group P21212 (cell constants *a*= 191.1 Å; *b*= 273.0 Å; *c*= 77.8 Å), whereas in the previous conditions the wt protein crystallized with DTT (PDB ID: 4UHW) exhibiting higher symmetry, in the tetragonal space group P42212 (cell constants *a*=*b*= 148.93 Å; *c*= 133.25 Å) (Coelho *et al*., 2015). The wt hAOX1 can also be crystallized with TCEP in the space group P21212 at pH 4.7 (data not shown).

**Figure 1.**
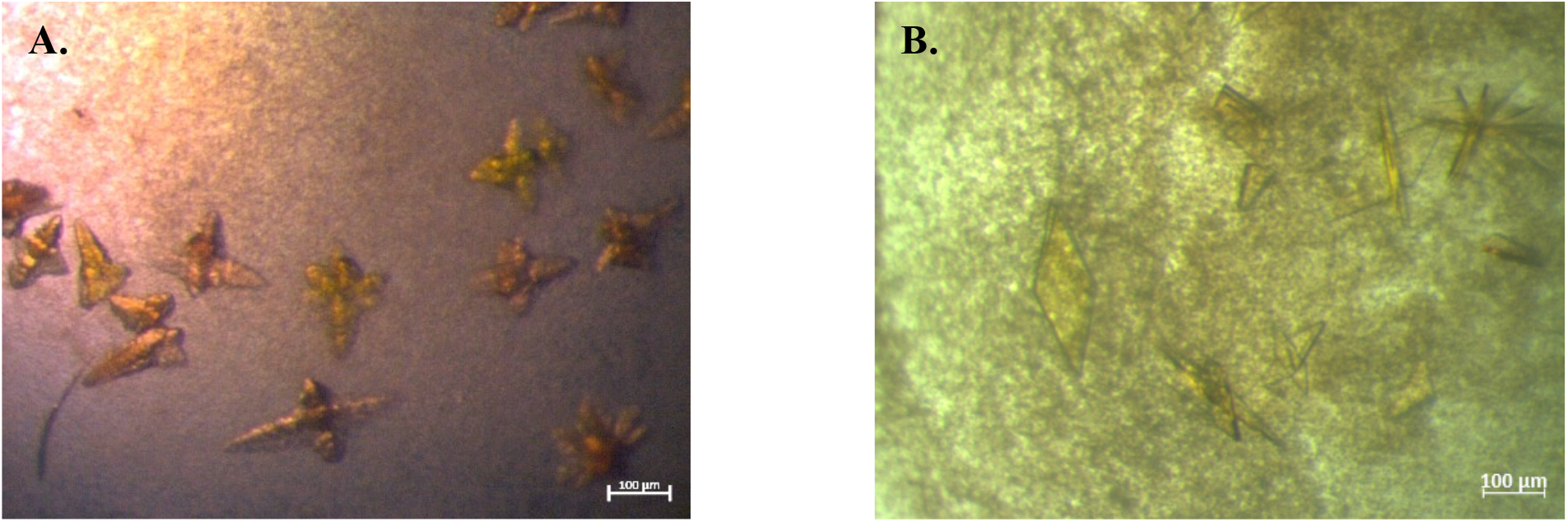
Images of hAOX1 crystals under the microscope. **(A)** Crystals formed in the presence of 30 mM DTT at pH 5.1, from the wt hAOX1. **(B)** Crystals formed upon the protein incubation with 1 mM TCEP at pH 4.7, observed for both wt and 6A variant.

**Figure 2.**
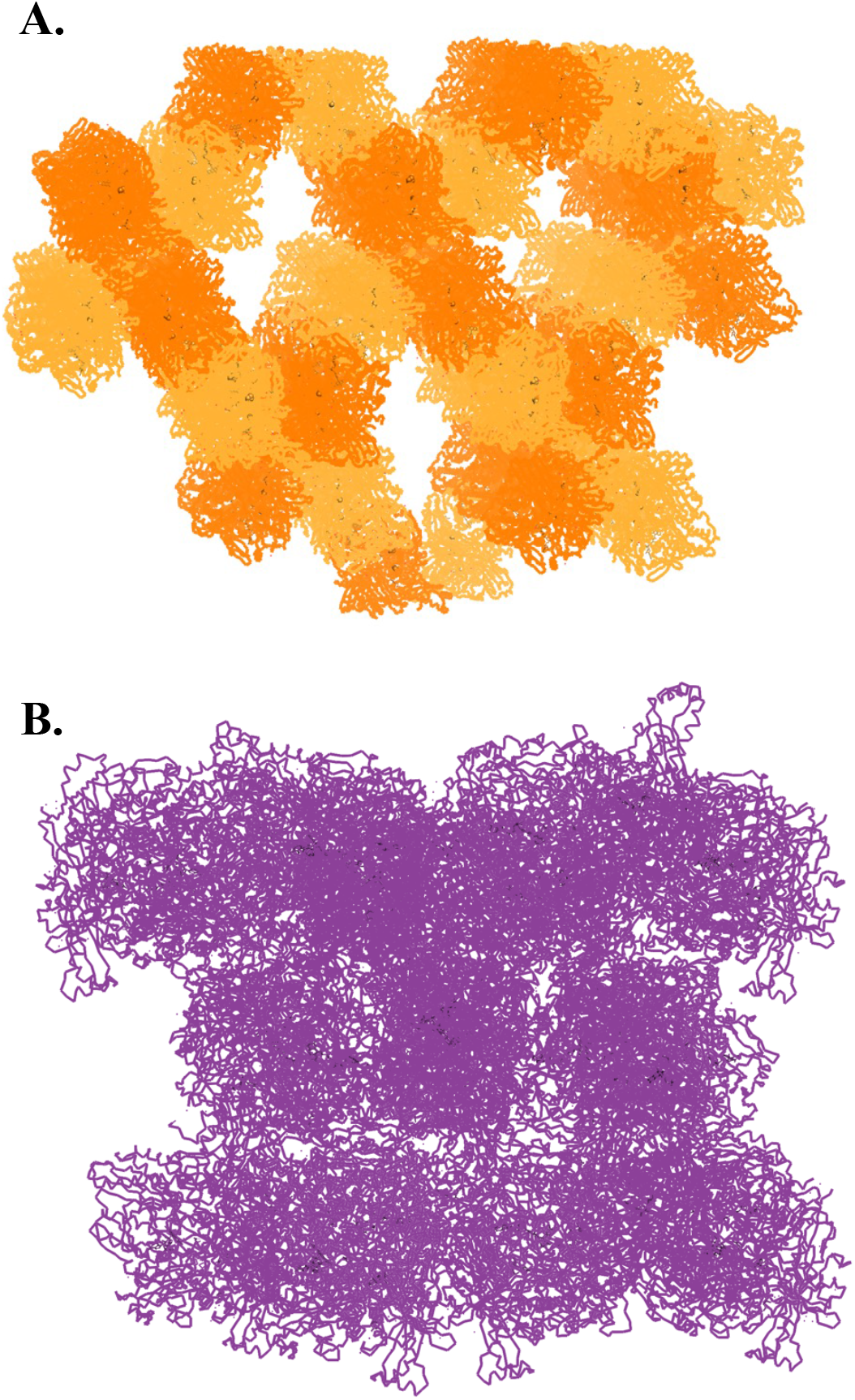
Difference in crystal packing for **(A) TCEP-grown** wt hAOX1 and hAOX1_6A at pH 4.7 and **(B)** DTT-grown wt hAOX1 (PDB:4UHW) at pH 5.1. Images created in PyMOL, by representing all symmetry-related molecules around a sphere with a radius of 100 Å (A) and 50 Å (B).

The two different crystal packing are depicted in Fig.2. The solvent content of TCEP-grown hAOX1_6A crystal is 63%, while for DTT-grown wt hAOX1 (PDB ID: 4UHW) is 50%. The asymmetric unit of this new crystal form is composed by two molecules forming the biological dimer through non-crystallographic symmetry. The overall structure of the biological dimer of TCEP-grown hAOX1_6A, as well as the shortest distances between each cofactor and their organization, is presented in Fig. 3.In DTT-grown wt hAOX1 crystals, only one molecule is present in the asymmetric unit, and the biological dimer is formed by a crystallographic dyad. After map interpretation, model building, and refinement, it was concluded that the overall fold of TCEP-grown hAOX1_6A is coincident to the DTT-grown wt hAOX1 with a Root Mean Square Deviation (RMSD) of 0.535 Å, for 2583 α-carbons (dimer). However, by superimposing chains A of DTT-grown wt hAOX1 (PDB ID: 4UHW) and TCEP-grown hAOX1_6A (RMSD 0.314 Å for 1138 α-carbons), the dimer shows a relaxed form with protomer B (chain B, RMSD 0.335 Å for 1167 α-carbons) in a slightly different conformation leading to increased distances between protomers, possibly as an internal rearrangement (Fig. 4). Domains II (FAD-containing domain) and III (Moco domain) are less superimposable compared to other regions, such as the dimerization interface. Chain B V540 is 3.0-Å apart from the same residue of the DTT-grown wt hAOX1. Analyzing the crystal structures, glutamate and aspartate surface-exposed residues are found to establish several crystalline intermolecular contacts. These interactions may be affected by changes in pH, leading to a different crystal packing and for example, side chains of residues E736, E958 and E1132 show slightly different orientations in each monomer (not shown).

**Figure 3.**
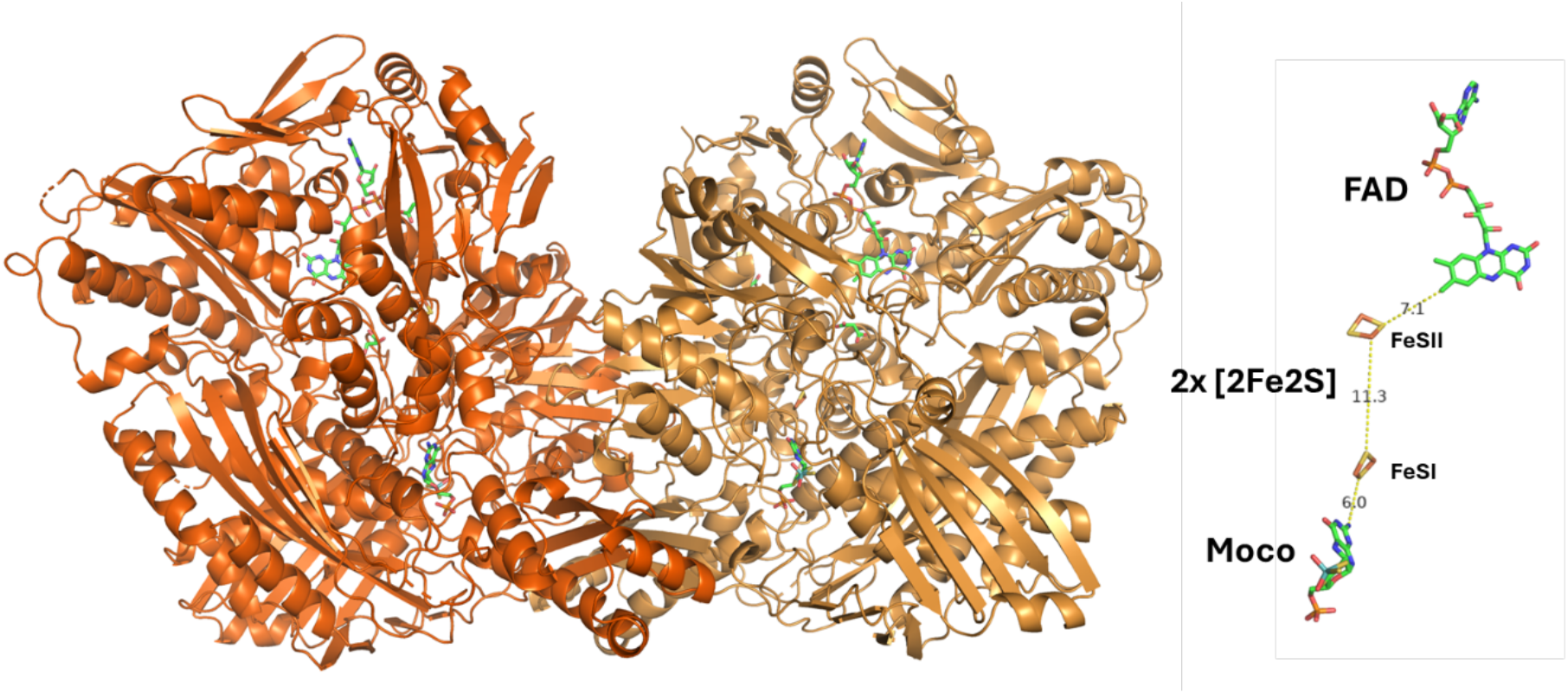
Homodimeric structure of hAOX1_6A (2×150 kDa) from crystals preincubated with TCEP in cartoon representation and on the right representation of the cofactors’ organization, as well as the shortest distances (in Å).

**Figure 4.**
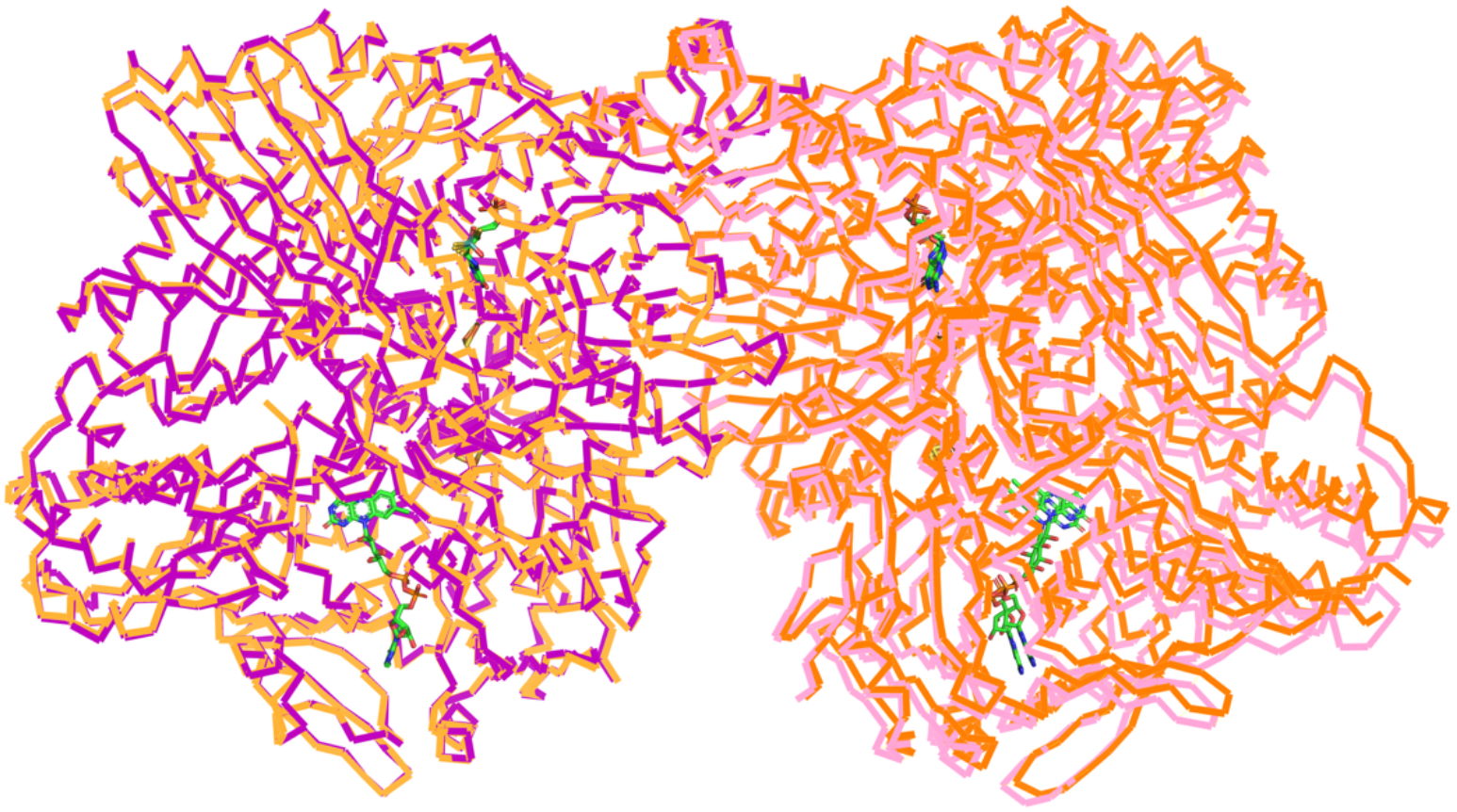
Superposition of hAOX1_6A in the P2_1_2_1_2 crystal form and hAOX1 in the P4_2_2_1_2 crystal form, by aligning chains A. The structures are represented as Ca, and each monomer is coloured differently but in the same shade. hAOX1_6A in the P2_1_2_1_2 in orange (chain A) and red (chain B), and wt hAOX1 in the P4_2_2_1_2 in purple (chain A) and magenta (chain B). The alignment of chains A (left monomers) show a RMSD of 0.350 Å, and the alignment of chains B (right monomers) present a RMSD of 0.372 Å.

The active site of the TCEP-grown hAOX1_6A, shown in Fig. 5, panel A, is similar to the DTT-grown wt enzyme being fully superimposable. The Mo coordination sphere contains two sulphurs from the pterin, as well as the sulphido (Mo=S) and hydroxyl (Mo-OH) groups at the equatorial plane and the oxo group (Mo=O) at the apical position. All the atoms from this center were refined with full occupancy and, in chain A, comparing with the amino acids from the vicinity (namely E1270, whose OE1 has a B-factor value of 47Å^2^), atoms from Moco have higher B-factors: The S1’ and S2’ from the pterin have 70 and 67 Å^2^, respectively; the Mo atom has 125 Å^2^, apical oxygen 125 Å^2^, equatorial oxygen 125 Å^2^ and the equatorial sulphur 123 Å^2^, suggesting that the centre is probably not fully occupied, but the sulphur atom was not removed upon the TCEP treatment, as the B-factor lies in the same order of magnitude as the atoms in the vicinity. The B-factors for this region in chain B are similar.

**Figure 5.**
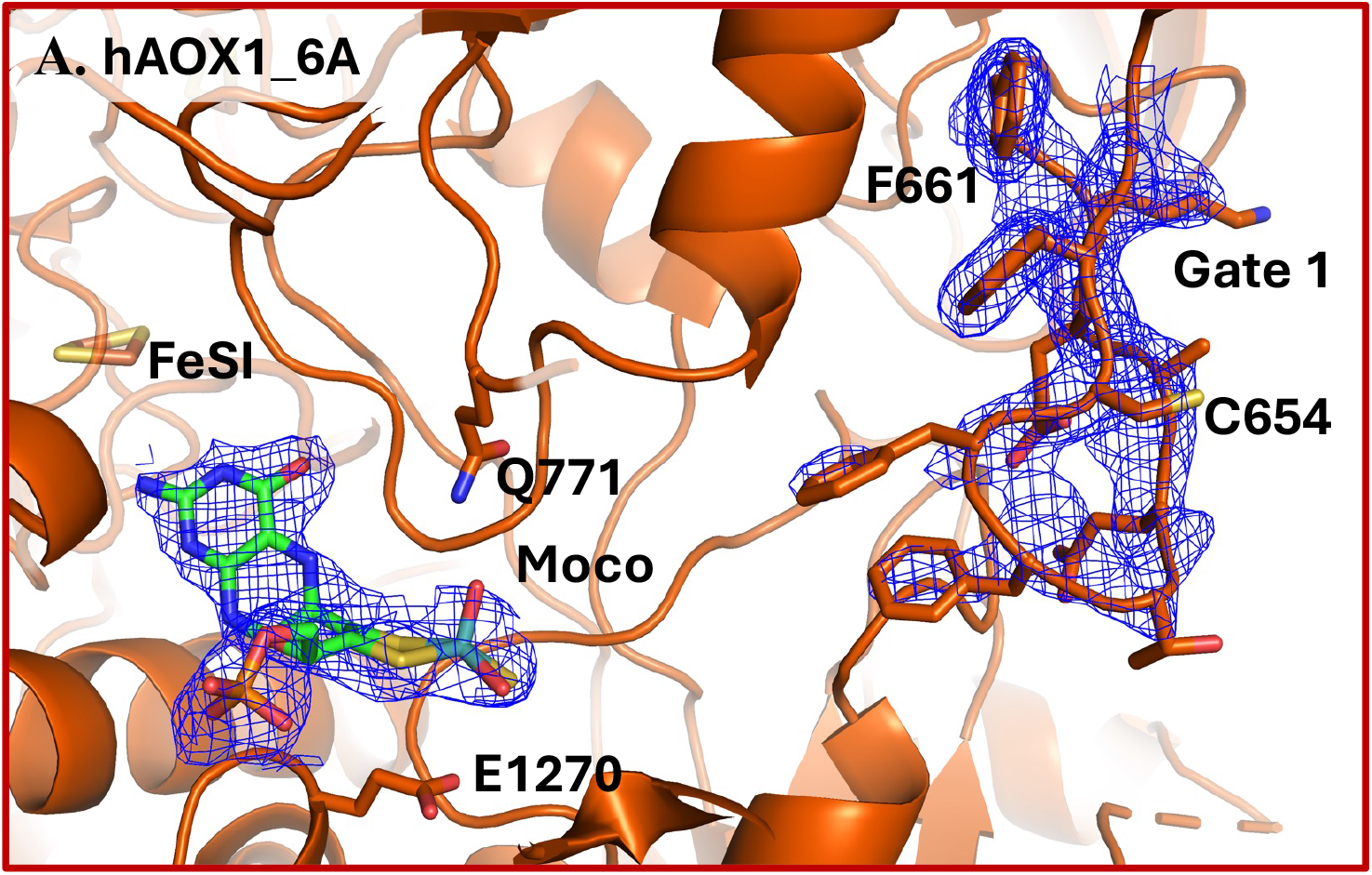
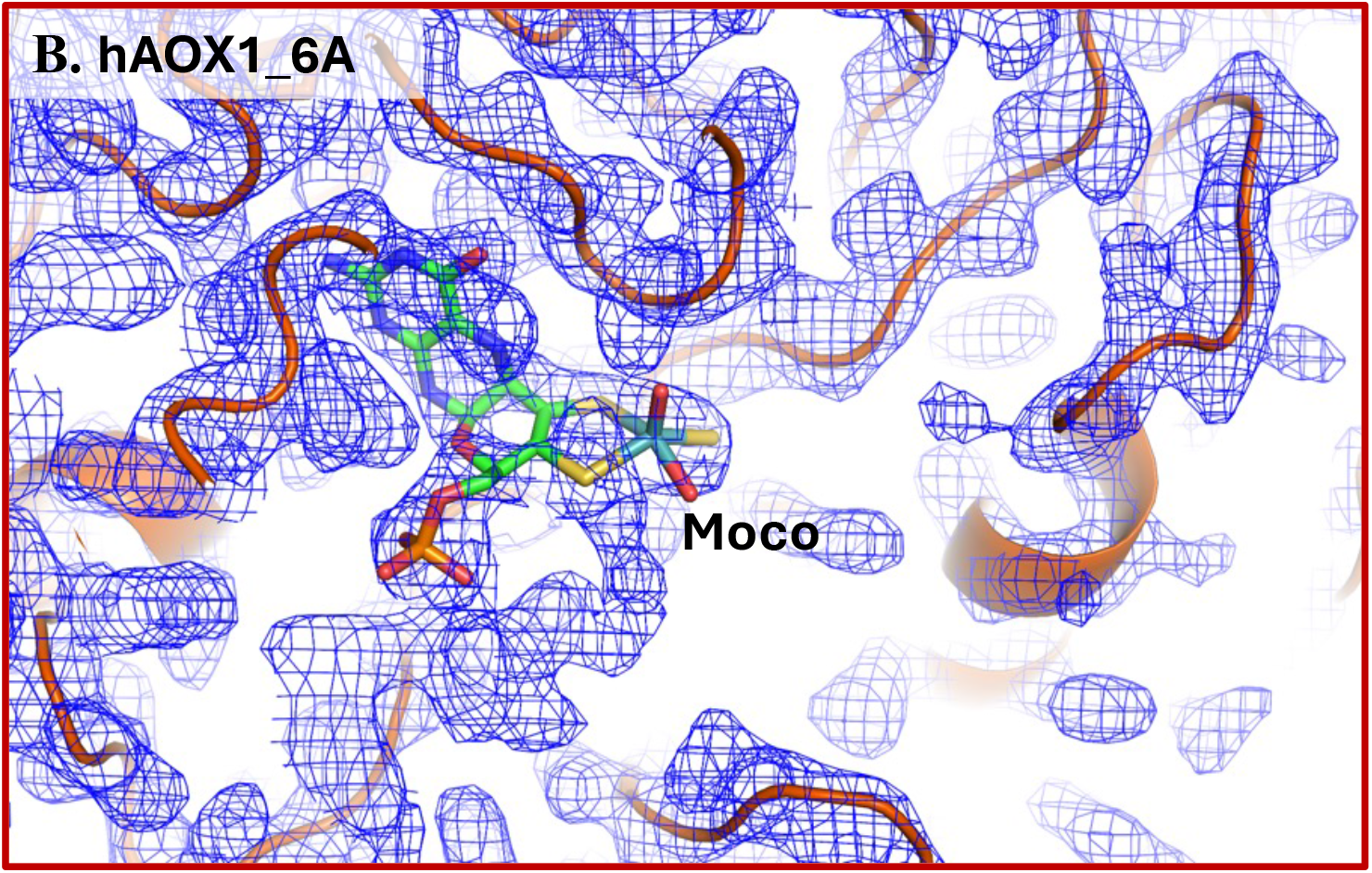
Electron density maps surrounding the active site of the free form of hAOX1_6A incubated with TCEP in two different orientations **(A)** Blue 2F_o_-F_c_ map (σ= 1.0) for Moco and Gate 1 (C_654_FFTEAEK_661_). **(B)** Blue 2F_o_-F_c_ map (σ= 1.0) of the vicinity of the Moco. Moco and FeSI cofactors represented by sticks and the main chain of the protein in dark orange, represented in cartoon. Images created in PyMOL.

Gate 1 (C654FFTEAEK661; near the substrate pocket entrance), which had no interpretable electron density in the previous models (DTT-grown wt hAOX1, PDB IDs: 4UHW and 4UHX), is now an ordered region and a model could be built (Fig. 5, panel A). This gate together with gate 2 is highly relevant for substrate and inhibitor recognition (Mota *et al*., 2019; Coelho *et al*., 2015). The vicinity of the active site (Fig. 5, panel B) shows the empty substrate pocket, as no additional electron density was observed during refinement cycles. No TCEP molecules are found neither in the vicinity of the Moco nor in the whole structure.

These data also allowed to model differently a solvent-exposed region containing the residues S1256QNSNT1261 in a new conformation compared to the DTT-prepared crystals (not shown) (Coelho *et al*., 2015; Mota *et al*., 2021). The solvent-exposed loop, (A160**C**KTF**C**KTSG**CC**QSKENGV**CC**…SQ231), with the mutated cysteines in bold, does not present any electron density for 22 residues (K167 to P189), underlining the flexibility of this loop. Nonetheless, the maps on the region C161A to C165A clearly show that the thiol groups are absent as expected, as shown negative electron density in Fo-Fc maps (Fig. 6). The final crystallographic statistics, regarding refinement, are shown in Table 4.

**Figure 6.**
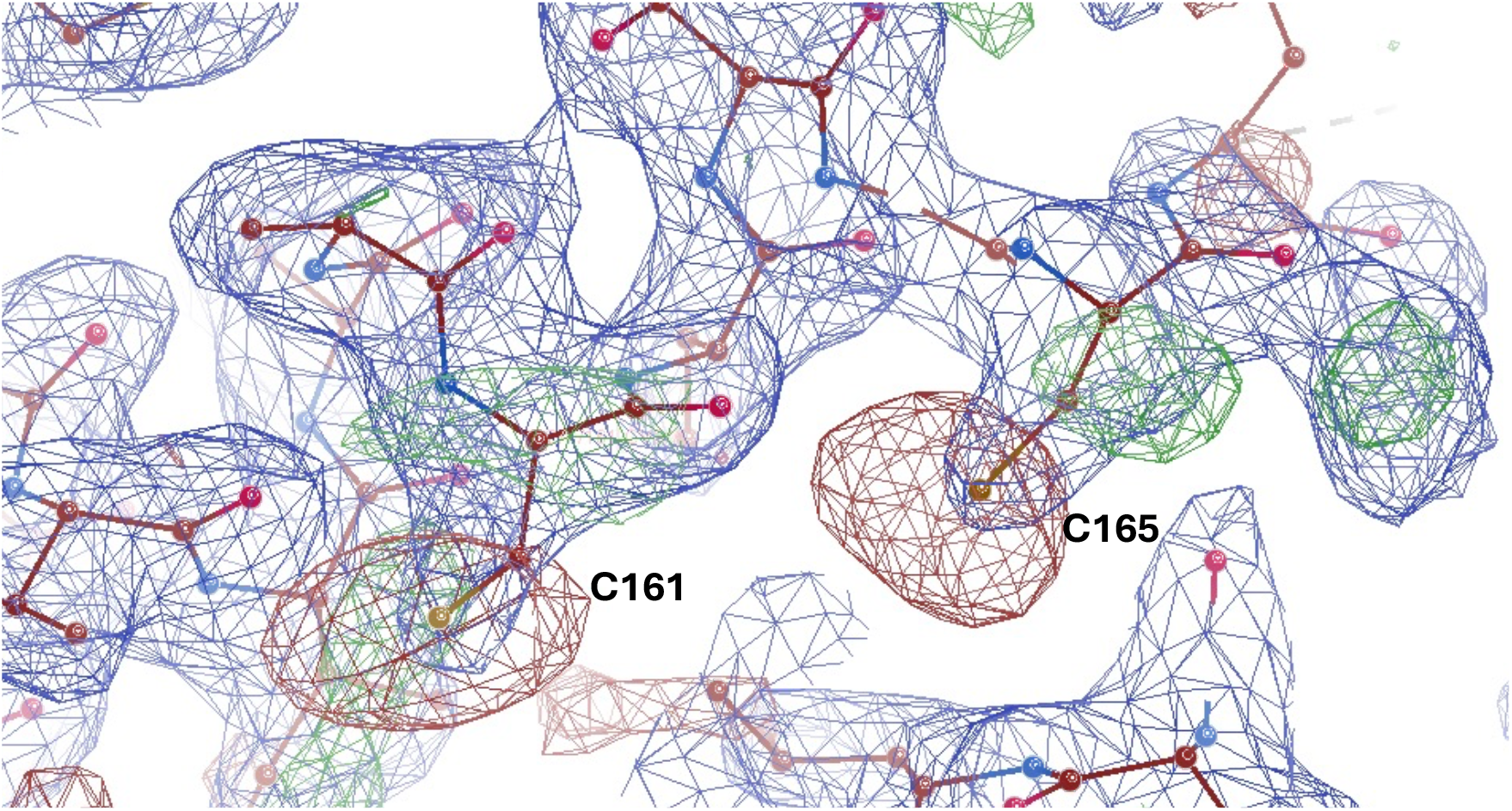
Electron density maps confirming the mutation of C161A and C165A on TCEP-grown hAOX1_6A. 2F_o_-F_c_ map (σ= 1.0) in blue and F_o_-F_c_ map (σ= 3.0) in green/red. In the refinement of the structure with C161 and C165 in the model, the sulfur atoms generate strong negative electron density (red). This finding confirms the C161A and C165A mutations in the hAOX1_6A construct.

### 3.2. hAOX1 activity assays with DTT and TCEP

Considering the inactivation of hAOX1 by DTT and the success of its crystallization in the presence of TCEP, it was analyzed if the use of TCEP as reducing agent would also inactivate the enzyme. The conditions described by Esmaeeli *et al*. were replicated with some adaptations (Esmaeeli *et al*., 2023). The wt hAOX1 was incubated with either DTT or TCEP for 30 minutes, 4 hours and O/N. The O/N incubation samples were desalted in a 10K centrifugal filter unit with buffer (50 mM Tris-HCl, 200 mM NaCl, 1mM EDTA, pH 8.0), to remove the reducing agents after a prolonged incubation and to assess whether the initial activity could be restored. In the kinetic assay phenanthridine (Phn) was used as substrate and the reaction started with the addition of the preincubated enzyme. The formation of the oxidized product (phenanthridinone, Pho, ε321=6400 M^-1^ cm^-1^) was followed at 321 nm. In Fig. 7 is possible to observe the relative initial reaction rate (v0), for each measurement. These values were obtained by calculating the slope in the linear part at the beginning of each kinetic curves and comparing each with the first control reaction (hAOX1 preincubated in buffer without any reducing agent for 30 minutes). It becomes evident that the incubation with DTT slows down the reaction, comparing to the treatment with TCEP or the controls. Over time, a clear cumulative effect of DTT became evident: after 30 minutes of incubation, the relative v_0_ decreased to 50%, after 4 hours it dropped to 11%, and following O/N incubation it reached approximately 10%. In the O/N assay with DTT, the activity was not recovered when hAOX1 was desalted. Upon incubation with TCEP, enzyme activity decreased approximately 40 %, comparing with the experiment with the highest v0 after 30 minutes and 4 hours, however, full activity was restored once TCEP was removed after O/N incubation (Fig. 7). Therefore, in agreement to a previous report (Esmaeeli *et al*., 2023), we observed that DTT inactivates hAOX1, without any activity recovering after the washing step. On the other hand, although TCEP lowers the enzyme activity (Fig. 7), upon its removal the enzyme regained its ability to oxidize the substrate with an activity similar to the control experiment (Fig. 7-O/N incubation and buffer exchange), demonstrating that TCEP is inhibiting hAOX1 in a reversible manner.

**Figure 7.**
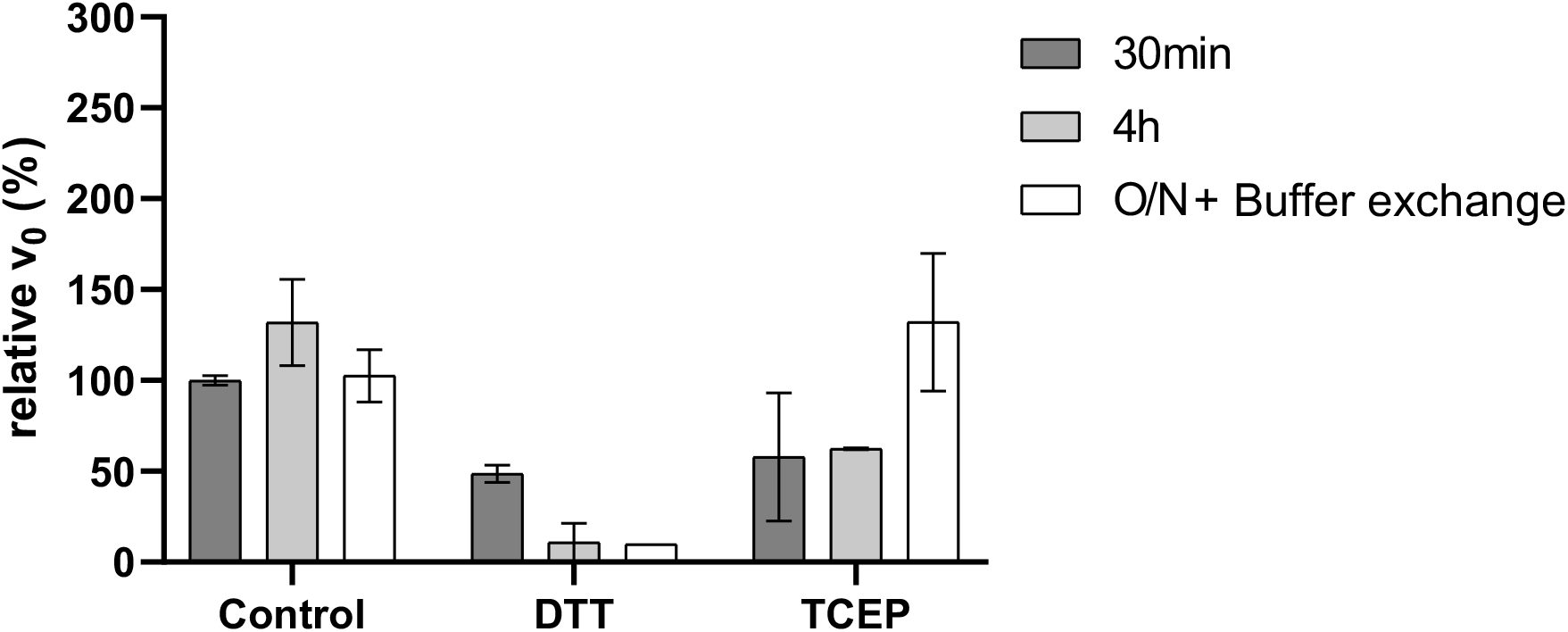
Relative initial reaction rate (v_0_) for each protein treatment. The values were normalized to the 30-minute incubation control (100%) with the standard deviation bars. The enzyme activity was measured after incubation with reducing agents (DTT or TCEP) for 30 minutes and 4 hours. After the O/N incubation, the enzyme was buffer exchanged to a buffer without reducing agents and activity measured. The control corresponds to wt hAOX1 without incubation with any reducing agent. The assay was performed in triplicates for each condition.

## 4. Conclusions

DTT has been widely used in studies of human aldehyde oxidase, however, it leads to enzyme inactivation. Its application in crystallization experiments may therefore compromise subsequent functional and structural studies. In this study, crystallization of both wt and the variant hAOX1_6A was achieved by replacing DTT with TCEP, a sulphur-free reducing agent. A new crystal form of hAOX1 is reported, revealing minor differences in the overall structure, as well as the absence of the reducing agent in the active site. Enzymatic assays showed that in the presence of TCEP, the wt hAOX1 has lower activity, but the reducing agent does not irreversibly inactivate the enzyme, in contrast to DTT.

Thus, the replacement of DTT with TCEP provides a practical strategy to circumvent enzyme inactivation during crystallographic studies. These findings facilitate future mechanistic investigations, including the trapping of catalytically relevant intermediates and the implementation of time-resolved crystallographic experiments.

## Acknowledgements

This work is financed by national funds from FCT—Fundação para a Ciência e a Tecnologia, I.P., through fellowship #NOVAID-B419 (to C.V.), in the scope of the project 2023.18077.ICDT; Research Unit Applied Molecular Biosciences—UCIBIO (UIDP/04378/2020 and UIDB/04378/2020) and Associate Laboratory Institute for Health and Bioeconomy—i4HB (LA/P/0140/2020). We acknowledge the European Synchrotron Radiation Facility for provision of synchrotron radiation facilities, and we would like to thank the staff of the ESRF, Grenoble for assistance and support in using beamline ID30A-3.

## Data availability

The atomic coordinates and structure factors of hAOX1_6A with TCEP are available in the Protein Data Bank (PDB) with the accession code 28MB.

